# A mitochondrial-immune axis drives the transcriptomic transition from brain aging to Alzheimer’s disease

**DOI:** 10.64898/2026.06.03.729900

**Authors:** Amita Pal, Shehbeel Arif, Ishanth Karthikeyan, Ethan Waisberg, Joseph W. Guarnieri

**Author notes:** Corresponding author. (J.W.G).

## Abstract

Aging is the primary risk factor for Alzheimer’s disease (AD), yet the molecular transitions linking normal brain aging to neurodegeneration remain poorly defined. Here, we performed integrative bulk transcriptomic analyses across a multi-region mouse aging atlas, a human aging-to-AD cohort, and an independent human AD validation dataset. Aging is associated with a progressive, region-specific increase in transcriptional perturbation, with the entorhinal cortex and choroid plexus showing the most pronounced age-associated remodeling. Females develop more extensive late-stage remodeling than males, characterized by stronger immune activation and greater suppression of mitochondrial metabolic pathways. Across cohorts, aging drives a coordinated shift toward immune activation and suppression of oxidative phosphorylation and respiratory-chain programs that is amplified in AD. Aged brains occupy an intermediate molecular state between young and AD conditions, supporting a continuum model. Together, our findings define a sex-modulated mitochondrial–immune axis linking normal aging to AD and highlight early immune–metabolic changes as potential intervention targets.

## INTRODUCTION

Alzheimer’s disease (AD) is the most common cause of dementia and a major global health burden (1–3). Although amyloid-β plaques and tau neurofibrillary tangles remain defining neuropathological features, it is increasingly clear that amyloid and tau pathology alone do not fully explain disease onset, regional vulnerability, clinical heterogeneity, or progression in sporadic AD (4–8). Recent biomarker and therapeutic advances have reinforced the importance of early disease staging, while also highlighting the need to understand downstream and parallel mechanisms, including neuroinflammation, metabolic failure, and loss of cellular resilience (9–13).

Aging is the strongest risk factor for AD, but the molecular processes by which normal aging becomes permissive for neurodegeneration remain incompletely understood (14–17). Transcriptomic and epigenomic studies indicate that normal brain aging is not uniform across regions or cell types; rather, aging produces spatially patterned changes involving glial activation, structural remodeling, altered chromatin states, and dysregulation of neuronal support programs (18–21). The entorhinal cortex (EC) is among the earliest cortical regions affected in AD (22,23), whereas the choroid plexus (CP), the principal cerebrospinal fluid-bluid barrier interface, undergoes substantial inflammatory remodeling during aging (24–27). Together, these observations suggest that age-associated molecular remodeling creates a vulnerable biological state upon which AD pathology is superimposed.

Mitochondrial metabolism is a central component of this vulnerability. Neurons have high energetic requirements and depend on oxidative phosphorylation (OXPHOS), tricarboxylic acid cycle activity, mitochondrial calcium buffering, mitochondrial trafficking, and mitophagy to sustain synaptic function (28–32). In AD, abnormalities in mitochondrial bioenergetics, respiratory-chain activity, reactive oxygen species generation, mitochondrial quality control, and mitochondrial DNA signaling have been reported early in disease progression (33–38). The mitochondrial cascade hypothesis posits that mitochondrial failure is not merely a late consequence of neurodegeneration but may actively contribute to disease initiation and amplification (30–33,35,39,40).

Neuroinflammation represents a second major axis of AD pathobiology. Genetic, transcriptomic and cellular studies have identified microglia and innate immune pathways as central regulators of AD risk and progression (41–50). Human AD microglia exhibit transcriptional programs that overlap with aging-related activation states while also displaying disease-specific features (51,52). Mitochondrial dysfunction and immune activation are mechanistically interconnected: impaired mitochondria promote inflammatory signaling through oxidative stress, altered metabolism, and release of mitochondrial damage-associated molecular patterns, whereas chronic inflammatory signaling further compromises mitochondrial function (53–63).

Sex is a critical but still under-resolved modifier of aging and AD biology. Women account for a disproportionate share of AD cases and often show distinct trajectories of risk, pathology, and clinical decline (64–68). Experimental and human studies suggest that sex hormones, sex chromosomes, immune regulation, and metabolic aging interact to shape female vulnerability (69–73). In particular, prior work has proposed that the aging female brain may enter a hypometabolic state earlier than the male brain, potentially lowering resilience to subsequent AD-related stressors (70,71).

Here, we integrated bulk transcriptomic analyses from a multi-region mouse aging atlas (19), a human aging-to-AD cohort (14,74), and an independent human AD validation dataset (52) to address four linked questions: whether normal aging produces region-specific transcriptional remodeling; whether these responses are sex-dependent; whether physiological aging occupies an intermediate molecular state between young healthy and AD brain; and whether mitochondrial and immune signatures are conserved across independent cohorts. Focusing on the EC and CP (regions of distinct relevance to cortical AD pathology and neuroimmune surveillance, respectively), we identify a sex-modulated mitochondrial-immune axis that links physiological aging to AD and is amplified during disease progression.

## RESULTS

### Aging induces progressive, region-specific transcriptional remodeling across the brain)

To define the transcriptional alterations associated with physiological brain aging, we analyzed bulk transcriptomic profiles across multiple brain regions, ages and sexes, using 3-month-old mice as the young reference baseline (**Fig. 1A**). We focused particularly on the EC and CP because the EC is among the earliest cortical regions affected in AD (22,23), whereas the CP serves as a critical neuroimmune and barrier-associated interface that undergoes substantial aging-related inflammatory remodeling(24–27).

**Figure 1.**
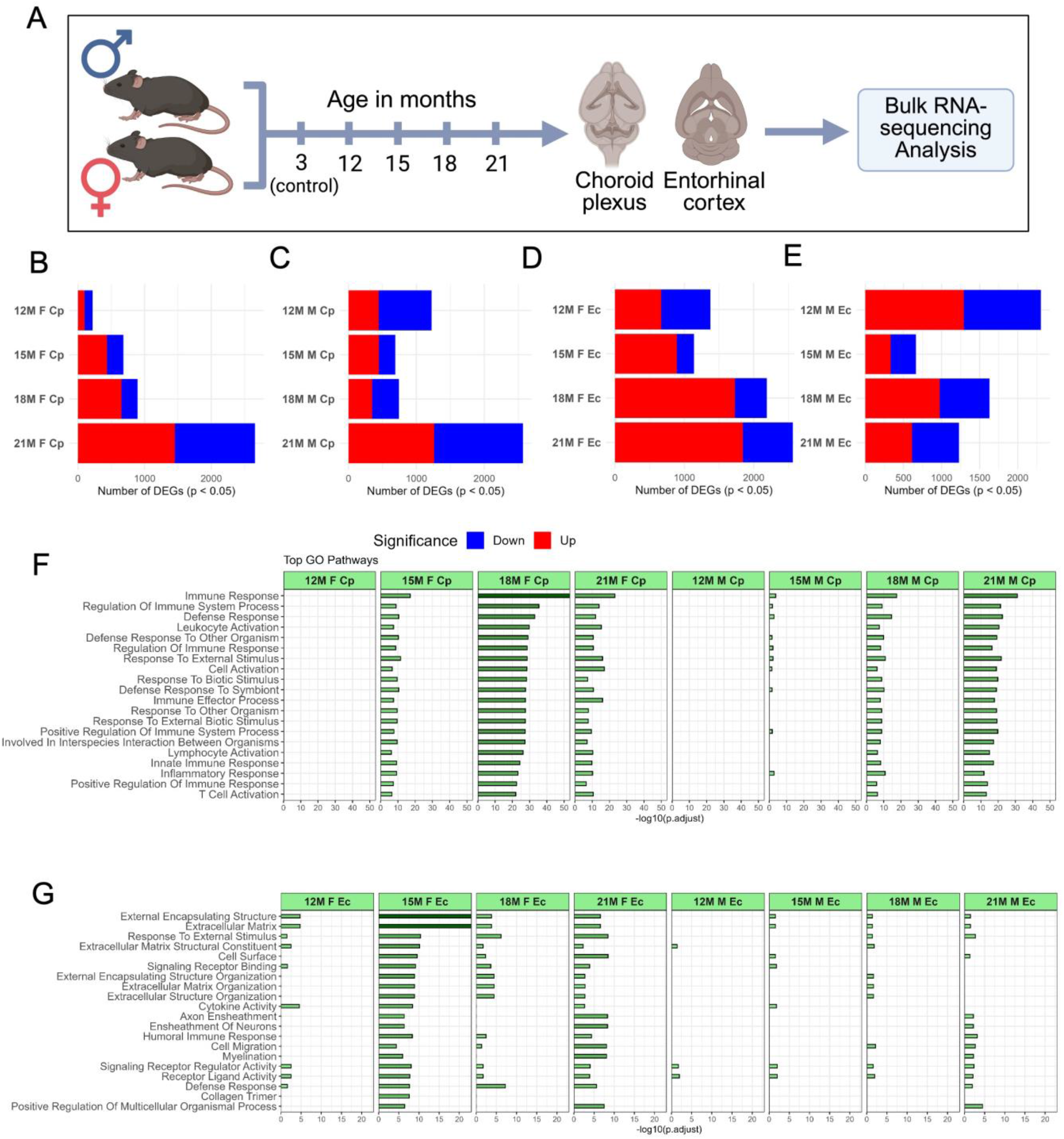
Aging drives progressive, region-specific and sex-dependent transcriptional remodeling across the brain. Bulk transcriptomes from multiple brain regions were profiled across aging time points in both sexes using 3-month-old mice as the young reference baseline. (A) Experimental schematic showing the mouse aging cohort (3, 12, 15, 18, and 21 months; both sexes) and the brain regions analyzed in this figure, including the entorhinal cortex (EC) and choroid plexus (CP). (B-E) Stacked bar plots showing the number of differentially expressed genes (DEGs) across aging time points and sex in the CP (B-C) and EC (D-E). Upregulated and downregulated DEGs (Benjamini–Hochberg adjusted *P* < 0.05) are shown in red and blue, respectively. Aging produces a progressive increase in transcriptional dysregulation with substantial regional and sex-dependent heterogeneity emerging at later stages. (F, G) Top enriched Gene Ontology (GO) Biological Process terms across the same CP (F) and EC (G) comparisons, with bar length representing –log_10_(adjusted *P*). EC aging is dominated by extracellular matrix organization and glial remodeling programs, whereas CP aging is dominated by immune and inflammatory pathways. Comparisons were performed against 3-month controls and stratified by sex.

Comparative analyses revealed a progressive increase in transcriptional dysregulation with advancing age across both EC and CP (**Fig. 1B-E**). The number of differentially expressed genes (DEGs) increased markedly between early and late aging stages, indicating progressive molecular destabilization. The magnitude and directionality of transcriptional remodeling, however, differed substantially between regions and sexes. Males exhibited a transiently higher number of DEGs during mid-aging stages (∼15 months), whereas females showed markedly greater transcriptional dysregulation at later ages, particularly at 21 months. This pattern suggests that males may undergo an earlier but comparatively limited adaptive transcriptional response, while females develop a more sustained and amplified late-aging remodeling program. Consistent with reported sex-dependent differences in neuroimmune regulation, hormonal decline, mitochondrial resilience and glial reactivity during aging (64–72).

Within the EC, females exhibited particularly strong transcriptional activation at later stages, with the 21-month female comparison showing extensive upregulation relative to young controls (1843 upregulated and 721 downregulated genes), indicative of robust induction of aging-associated transcriptional programs. In contrast, male EC samples showed comparatively lower DEG burdens and more balanced transcriptional changes across late aging stages. Within the CP, aging-associated transcriptional changes emerged progressively with age in both sexes, although overall DEG distributions remained more moderate than those observed in female EC samples. Expanded volcano plots for all EC and CP comparisons are provided in **Supplementary Fig. 1**.

Volcano plot analyses further demonstrated age-dependent increases in both the magnitude and statistical significance of transcriptional alterations across EC and CP comparisons (**Fig. S1**). Early aging stages exhibited relatively compact DEG distributions, whereas later stages showed broader transcriptional dispersion and larger fold-change magnitudes, consistent with progressive molecular remodeling during aging (18–20). Several late-stage EC comparisons displayed marked asymmetry between upregulated and downregulated genes, indicative of activation-dominant transcriptional responses in vulnerable cortical regions.

Region-specific Gene Ontology (GO) analyses identified distinct molecular trajectories between EC and CP. Aging in the CP was characterized predominantly by enrichment of immune-associated pathways, including cytokine signaling, leukocyte activation, chemotaxis and inflammatory-response programs, consistent with its established role as a neuroimmune signaling interface (24–27). In contrast, the EC showed stronger enrichment of extracellular matrix organization, glial remodeling and structural reorganization pathways, suggesting progressive disruption of neuronal and tissue homeostasis during aging (19,20) (**Fig. 1F**,**G**).

### SEX-dependent aging responses reveal enhanced immune activation and mitochondrial metabolic suppression in females

To determine whether sex modifies aging-associated molecular remodeling, we performed sex-stratified pathway analyses across aging time points and brain regions. Females consistently exhibited stronger transcriptional perturbation than males, particularly within immune-associated and mitochondrial pathways (**Fig S1**).

Pathway-level heatmap analyses revealed widespread enrichment of inflammatory signaling programs in females, including cytokine-mediated signaling, complement activation, leukocyte activation, adaptive immune response and microglial-associated pathways (**Fig 2B**). These immune signatures became progressively stronger with advancing age, especially within EC and CP samples. In parallel, females showed more pronounced suppression of mitochondrial metabolic pathways than males. Pathways associated with OXPHOS, ATP synthesis, respiratory electron transport, mitochondrial translation, tricarboxylic acid (TCA) cycle metabolism and mitochondrial complex assembly were consistently downregulated in aged female brains. These alterations were particularly evident in EC samples at later aging stages, indicating enhanced mitochondrial vulnerability in females (67,70,72,75,76).

**Figure 2.**
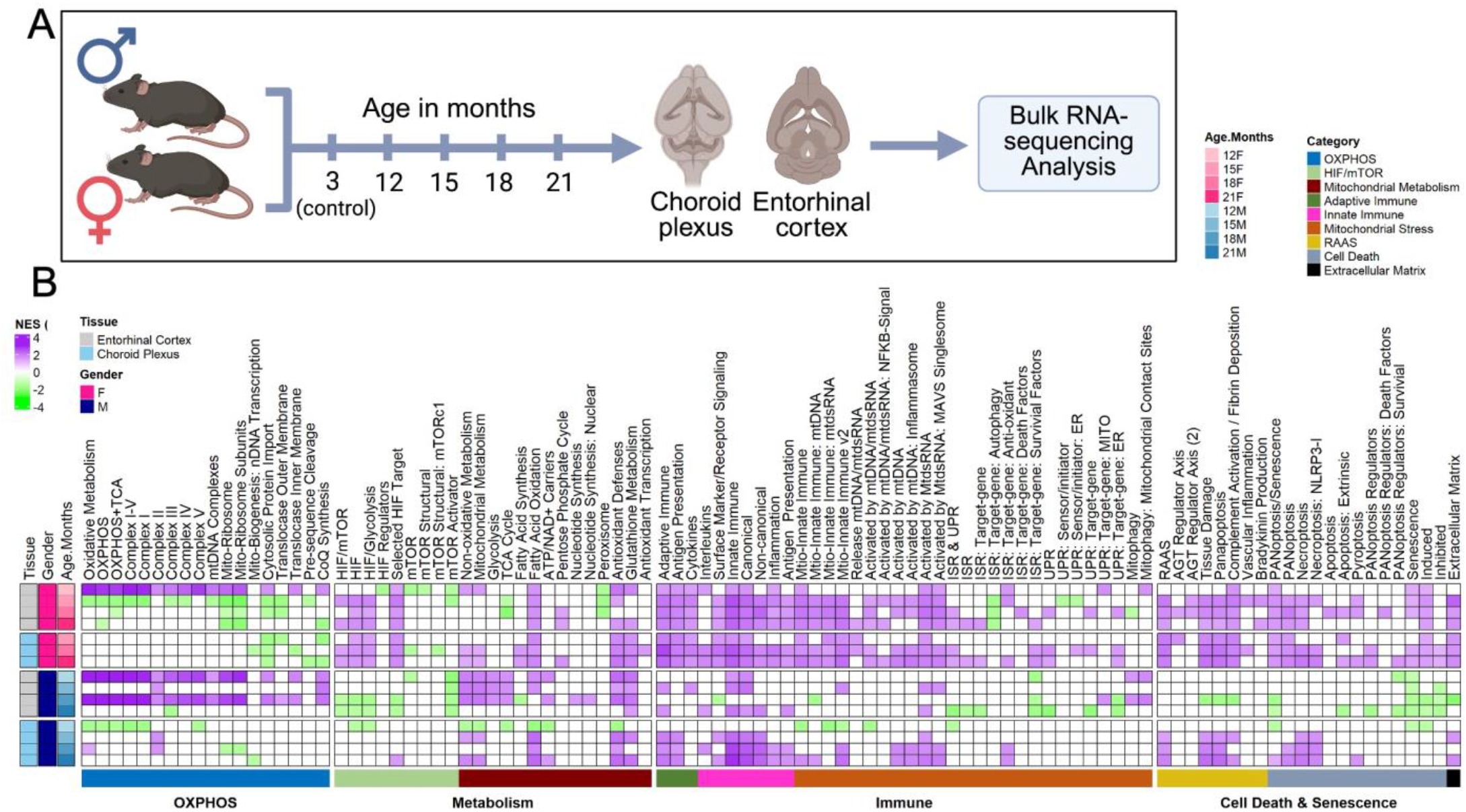
Female aging is associated with enhanced immune activation and stronger suppression of mitochondrial metabolic pathways. Sex-stratified pathway enrichment across aging time points and brain regions reveals a coordinated, female-biased immune–metabolic axis. (**A)** Experimental schematic for the mouse aging cohort (3, 12, 15, 18, and 21 months; both sexes) and brain regions analyzed (EC, CP). (**B)** Heatmap of normalized enrichment scores (NES) from gene set enrichment analysis (GSEA) across curated pathway collections, organized into major thematic categories: OXPHOS (oxidative phosphorylation, mitochondrial ribosome, mtDNA-encoded subunits, ETC assembly), mitochondrial and non-oxidative metabolism (TCA cycle, fatty acid oxidation, nucleotide synthesis, antioxidant defenses), HIF/mTOR signaling, adaptive and innate immune programs (cytokines, antigen presentation, inflammasome, NF-κB), mtDNA-driven innate immune signaling, integrated stress response and unfolded protein response, RAAS signaling, PANoptosis/senescence and extracellular matrix organization. Rows correspond to individual comparisons annotated by tissue (EC, CP), sex (F, M) and age (12, 15, 18, 21 months) relative to 3-month controls. Female samples consistently show stronger positive enrichment of immune programs (purple) together with stronger negative enrichment of mitochondrial and OXPHOS programs (green) than male samples, with these signatures intensifying at later aging stages. Enrichment significance: Benjamini– Hochberg adjusted *P* < 0.05.

At the gene level, female aging was associated with stronger induction of inflammatory and microglial-associated transcripts including *Trem2, Tyrobp, C1qa, C1qb* and *Cd68* (41,42,46–49,77,78). Conversely, mitochondrial respiratory-chain genes encoding components of all the electron transport chain complexes, showed greater suppression in females than in males (**Fig S1**). These gene-level changes closely paralleled pathway-enrichment analyses and collectively support the emergence of a sex-dependent immune–metabolic aging axis.

Together, these findings indicate that female brain aging is characterized by enhanced inflammatory activation coupled with stronger mitochondrial metabolic suppression. This coordinated immune–metabolic remodeling may contribute to the increased susceptibility of females to age-associated neurodegenerative processes (64,66,68,69).

### Aging represents an intermediate molecular state between healthy and Alzheimer’s disease brain

To investigate the molecular relationship between physiological brain aging and AD, we comparatively analyzed transcriptomic profiles from young controls, aged controls, and AD brain samples (79–81). Differential expression analyses revealed progressive transcriptional dysregulation across the aging-to-AD continuum, with the highest DEG burden observed in aged AD samples relative to young controls (**Fig. 3B**). In contrast, aged-versus-young control comparisons exhibited comparatively modest transcriptional remodeling, indicating that physiological aging induces selective molecular alterations that become amplified during AD progression (14,74,82).

**Figure 3.**
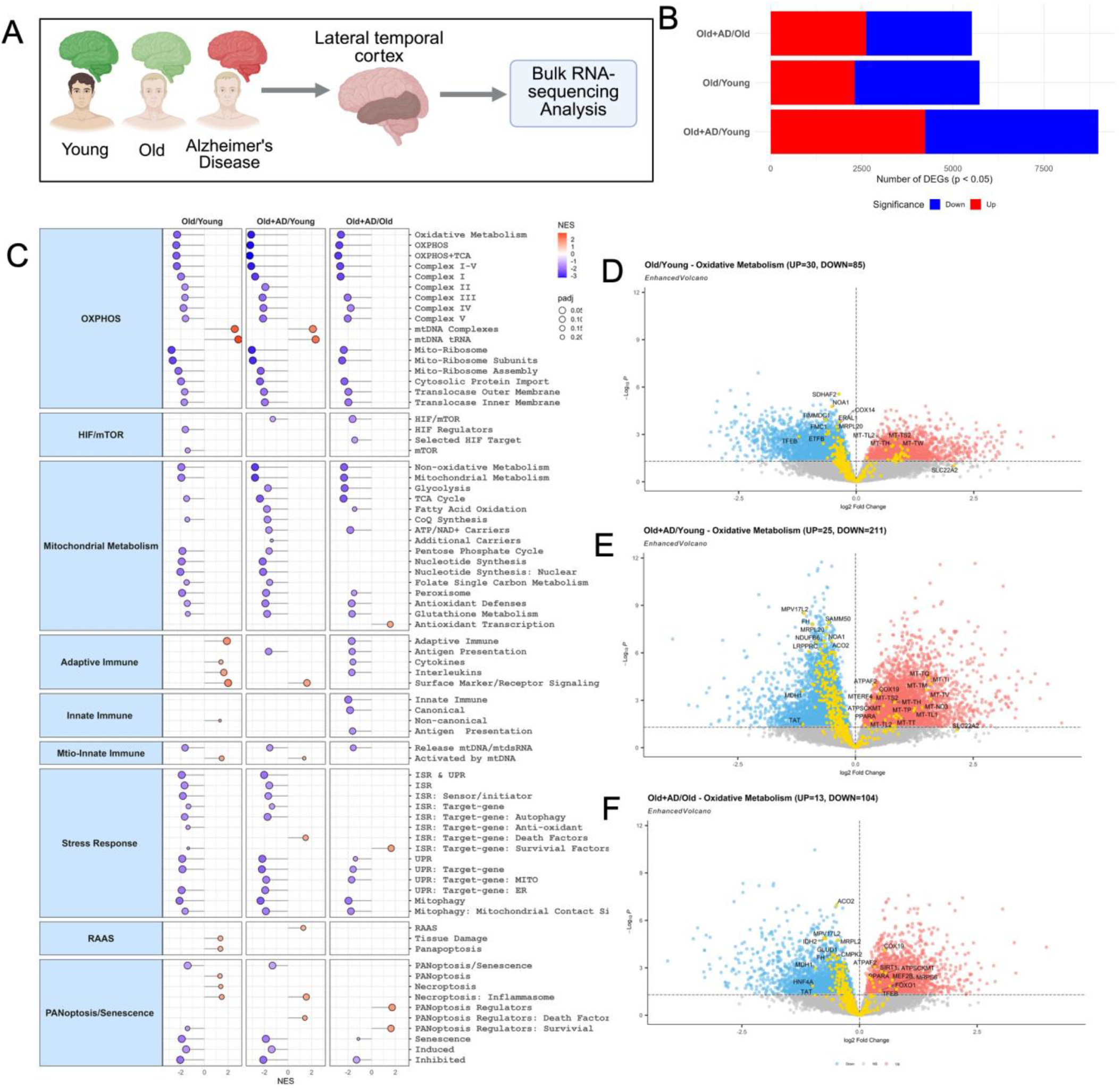
Aging occupies an intermediate molecular state between young healthy and Alzheimer’s disease brain. Integrative comparison of young healthy, aged healthy and AD human brain transcriptomes revealing progressive immune activation and mitochondrial metabolic suppression along the aging-to-AD continuum. (A) Schematic summary of the integrative aging-to-AD comparison framework (Old/Young, Old+AD/Young and Old+AD/Old). (B) Stacked bar plots showing the number of DEGs (adjusted *P* < 0.05) for each pairwise comparison; upregulated and downregulated genes are shown in red and blue, respectively. DEG burden is highest in the Old+AD versus Young comparison. c GSEA dot plot showing NES across curated pathway collections (OXPHOS, HIF/mTOR, mitochondrial metabolism, adaptive and innate immune, mitochondria-driven innate immune, ISR/UPR, RAAS, PANoptosis/senescence) for the three comparisons. Dot color encodes NES and dot size reflects adjusted *P* value; immune pathways become progressively activated and mitochondrial pathways progressively suppressed from Old/Young to Old+AD/Young. (D-F) Oxidative-metabolism–focused volcano plots for Old/Young (D), Old+AD/Young (E) and Old+AD/Old (F). Each point represents a gene from the curated oxidative metabolism gene set, plotted by log_2_ fold change against –log_10_(adjusted *P*). Genes passing adjusted *P* < 0.05 are colored by direction (red, up; blue, down); yellow points pass only the effect-size threshold (|log_2_ fold change| ≥ 0.5); grey points are not significant. UP and DOWN counts are indicated above each panel. Selected top genes are labeled.

Pathway-enrichment analyses revealed progressive activation of immune and stress-associated pathways alongside suppression of mitochondrial and oxidative metabolic programs across the transition from young control to aged and AD brain (**Fig. 3C**). Immune-related pathways, including cytokine signaling and innate immune activation, showed increasing enrichment in AD-associated comparisons, whereas oxidative phosphorylation, ATP biosynthesis and respiratory electron transport pathways exhibited progressive decline (28,28,30–32,34,35,38,83).

Oxidative metabolism–focused volcano plots further demonstrated stage-dependent metabolic dysregulation (**Fig. 3D-F**). The aged-versus-young control comparison showed relatively limited perturbation of oxidative metabolism genes, whereas the aged AD-versus-young comparison displayed substantially broader transcriptional dispersion and marked dysregulation of mitochondrial and respiratory metabolism–associated genes. Although oxidative metabolic alterations remained detectable in the aged AD-versus-aged comparison, the magnitude of these changes was comparatively reduced, suggesting that components of mitochondrial dysfunction emerge during aging and are subsequently exacerbated in AD (30–32,84).

Comparison-specific GO analyses supported these findings (**Fig. S2A-C**). Physiological aging was associated predominantly with pathways related to myelination, gliogenesis and structural remodeling (19,20), whereas aged AD samples showed strong enrichment of mitochondrial metabolic processes, including ATP metabolism, aerobic respiration and proton transmembrane transport. Enhanced volcano plots further confirmed progressively increased transcriptional dispersion and molecular dysregulation in AD-associated comparisons (**Fig. S2D-F**).

Together, these findings support a model in which physiological aging represents an intermediate molecular state characterized by mild inflammatory activation and early metabolic adaptation, whereas AD is associated with amplified neuroimmune activation and pronounced mitochondrial metabolic dysfunction (79,81,84).

### Mitochondrial metabolic dysfunction and immune activation are conserved across independent Alzheimer’s disease datasets

To validate the reproducibility of the mitochondrial–immune signatures identified in our primary analyses, we analyzed an independent AD transcriptomic cohort derived from fusiform gyrus tissue [Fig. 4). We compared male and female subjects under non-AD and AD conditions to determine whether sex-dependent transcriptional differences observed during aging are altered in Alzheimer’s disease. Cross-cohort comparisons revealed highly consistent molecular alterations characterized by coordinated inflammatory activation and suppression of mitochondrial metabolic pathways, supporting the presence of a conserved mitochondrial– immune axis in AD-associated neurodegeneration (53–58).

**Figure 4.**
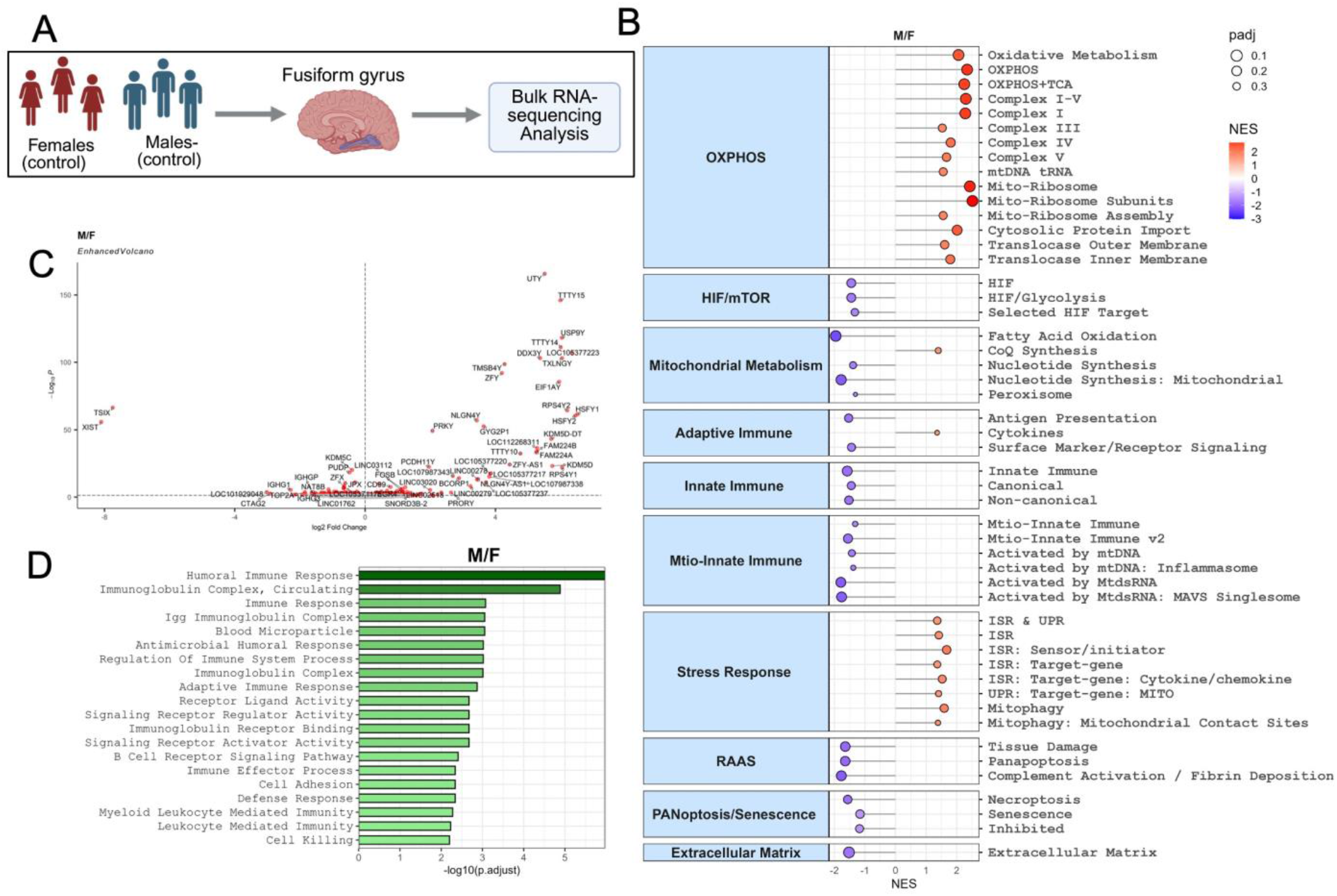
Mitochondrial metabolic suppression and immune activation are conserved across independent Alzheimer’s disease datasets. Validation in an independent human AD transcriptomic dataset (fusiform gyrus) confirms a conserved mitochondrial–immune axis. **(A)** Schematic summary of the validation framework comparing aged (non-AD) male versus female (M/F). **(B)** GSEA dot plot of NES across the curated pathway collections used in Fig. 2, shown for the M/F, AD M and AD F comparisons. Dot color encodes NES and dot size reflects adjusted *P*. AD samples of both sexes show coordinated activation of inflammatory, immune and stress-associated pathways together with suppression of OXPHOS, ATP synthesis and mitochondrial respiratory-chain programs. **(C)** Enhanced volcano plot for the M/F comparison, highlighting sex-chromosome–encoded transcripts as a positive technical control. (**D)** Bar plots of the top 20 enriched GO Biological Process terms for the M/F. Bar length represents –log_10_(adjusted *P*). Enriched pathways are predominantly associated with humoral immunity, immunoglobulin-mediated responses, adaptive immune signaling, leukocyte-mediated immunity, receptor signaling, and immune effector functions.

In age-matched control subjects, male-versus-female analysis revealed a distinct sex-associated transcriptional signature characterized by differential enrichment of mitochondrial, metabolic, immune, stress-response, and extracellular matrix pathways (**Fig. 4B-C**). Males showed stronger enrichment of oxidative phosphorylation and mitochondrial metabolic pathways, including OXPHOS, electron transport chain complexes, mitochondrial translation, and mitochondrial protein import signatures. In contrast, females showed relatively greater enrichment of immune-associated pathways, including adaptive immune signaling, innate immune activation, cytokine signaling, antigen presentation, and mito-innate immune pathways. Stress-response programs, including ISR/UPR-related pathways, mitophagy, and inflammatory remodeling signatures, were also more prominent in females (**Fig. 4B**). Gene ontology analysis further supported this sex-dependent immune bias, with enrichment of humoral immune response, immunoglobulin complex, receptor signaling, leukocyte -mediated immunity, and cell-killing pathways in females (**Fig. 4D**).

We next asked whether these sex-dependent differences were preserved or amplified in AD subjects from the same dataset. Differential expression analyses revealed substantial transcriptional perturbation in AD brains relative to controls across both sexes (**Fig. 5B**). Although male AD samples exhibited a modestly greater overall DEG burden, females showed stronger enrichment of inflammatory and mitochondrial metabolic pathways. These findings suggest that male brains may undergo broader but more heterogeneous transcriptional remodeling, whereas female brains exhibit more coordinated disease-relevant immune–metabolic dysregulation (64,65,68,69).

**Figure 5.**
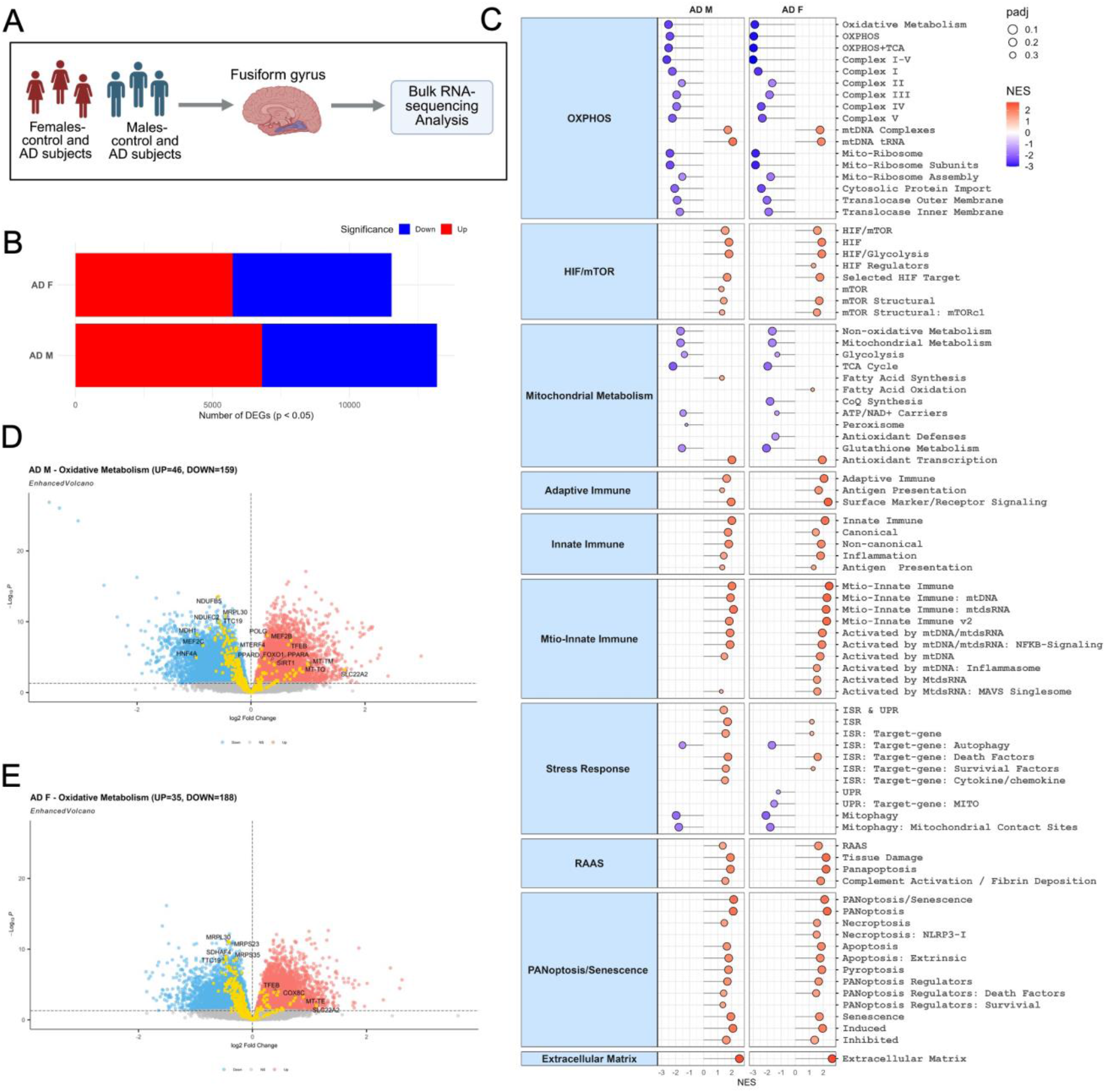
Mitochondrial metabolic suppression and immune activation are conserved across independent Alzheimer’s disease datasets. Validation in an independent human AD transcriptomic dataset (fusiform gyrus) confirms a conserved mitochondrial–immune axis. **(A)** Schematic summary of the validation framework comparing male versus female (M/F), AD female versus female control (AD F) and AD male versus male control (AD M). **(B)** Stacked bar plots showing the number of DEGs (adjusted *P* < 0.05) for each comparison; upregulated and downregulated genes are shown in red and blue, respectively. AD samples of both sexes show extensive transcriptional perturbation relative to controls. **(C)** GSEA dot plot of NES across the curated pathway collections used in Fig. 2, shown for the AD M and AD F comparisons. Dot color encodes NES and dot size reflects adjusted *P*. AD samples of both sexes show coordinated activation of inflammatory, immune and stress-associated pathways together with suppression of OXPHOS, ATP synthesis and mitochondrial respiratory-chain programs. (**D-E)** Oxidative-metabolism–focused volcano plots for AD M (**D**) and AD F (**E**). Each point represents a gene from the curated oxidative metabolism set, plotted by log_2_ fold change against –log_10_(adjusted *P*). Coloring as in Fig. 3D–F. AD samples of both sexes show coordinated suppression of mitochondrial respiratory-chain and ATP-synthesis genes together with induction of inflammatory and glial-associated transcripts; UP and DOWN counts are indicated above each panel.

Pathway enrichment analyses demonstrated robust activation of inflammatory and immune-associated programs in AD brains, including cytokine signaling, leukocyte activation, adaptive immune responses, NF-κB signaling, and stress-associated pathways (**Fig. 5C**) (43,45,50,85). Concurrently, pathways associated with oxidative phosphorylation, ATP biosynthesis, respiratory electron transport, and mitochondrial translation were consistently suppressed, indicating widespread disruption of neuronal bioenergetic homeostasis (28,29,35,83,86–89).

Oxidative-metabolism–focused volcano plot analyses further refined these observations (**Fig. 5D-E**). AD-associated comparisons revealed coordinated suppression of mitochondrial respiratory-chain and oxidative phosphorylation genes, including *NDUFS1, NDUFA9, SDHB, UQCRC1, COX5A*, and *ATP5F1A*, consistent with impaired electron transport chain activity and reduced mitochondrial ATP production (34,83,86,90). In parallel, inflammatory and glial-associated transcripts, including *TREM2, TYROBP, C1QA, C1QB, CD68*, and *GFAP*, were significantly upregulated, indicating persistent neuroimmune activation (41–43,45–48,51,52,85,91) (**Fig. S3C-D**)., These gene-level alterations closely aligned with the pathway enrichment results, demonstrating that oxidative metabolic dysfunction in AD reflects coordinated suppression of broader mitochondrial bioenergetic programs rather than isolated transcriptional changes.

GO pathway analyses further showed persistent enrichment of synaptic signaling, receptor activity and trans-synaptic communication pathways, while immune-regulatory and leukocyte-associated pathways became increasingly prominent in AD samples (**Fig. S3A-B**).

Together, these findings indicate that sex differences in the aged human brain are already evident in the absence of AD, with females showing stronger immune and stress-response signatures and males showing relatively stronger mitochondrial oxidative metabolism. In AD, this baseline sex divergence becomes more pronounced, with females displaying enhanced immune activation, mitochondrial-immune signaling, ISR/UPR engagement, and inflammatory cell-death-associated transcriptional programs. These results suggest that AD amplifies pre-existing sex-dependent mitochondrial–immune differences in the aged human brain. (53–58,92– 94).

## DISCUSSION

Here, we identify a sex-modulated mitochondrial–immune axis that links physiological brain aging to AD. Across independent mouse aging and human AD transcriptomic cohorts, progressive aging was associated with coordinated immune activation, suppression of mitochondrial metabolic pathways, and pronounced regional and sex-dependent heterogeneity. These molecular signatures were substantially amplified in AD, supporting a continuum model in which aging and neurodegeneration share a common biological trajectory rather than representing entirely distinct processes (79–81).

A central finding of this study is the consistent coupling between inflammatory activation and mitochondrial dysfunction. Both aging and AD datasets revealed coordinated suppression of oxidative phosphorylation, respiratory electron transport, ATP synthesis, and mitochondrial translation pathways together with enrichment of inflammatory and microglial-associated programs. Oxidative metabolism–focused volcano analyses further revealed progressive dysregulation of mitochondrial respiratory-chain genes during the transition from physiological aging to AD. These observations align with previous studies implicating mitochondrial dysfunction as a major contributor to neuronal vulnerability and AD pathogenesis (34,38,83,86). Given the high energetic demands of neurons, even moderate age-associated reductions in mitochondrial function may impair synaptic maintenance, proteostasis, and cellular stress responses, thereby lowering resilience to neurodegenerative stressors (31,33,87).

Concurrently, immune-associated pathways including cytokine signaling, complement activation, leukocyte activation, and microglial programs were progressively enriched during aging and markedly amplified in AD. Increased expression of inflammatory transcripts including *TREM2, TYROBP, C1QA, C1QB, CD68*, and *GFAP* supports sustained activation of innate immune and glial responses (46–48,77,78,95,96). The reciprocal relationship between mitochondrial dysfunction and inflammation suggests the presence of a reinforcing immune–metabolic feedback loop during aging and neurodegeneration (56,58). Activated glial states can alter tissue metabolism and increase oxidative stress, whereas mitochondrial dysfunction itself promotes inflammatory signaling through reactive oxygen species generation, impaired mitophagy, and release of mitochondrial danger-associated signals (53–55,57).

Our analyses further reveal substantial regional heterogeneity in aging-associated remodeling. The entorhinal cortex exhibited extensive late-stage transcriptional dysregulation characterized by immune activation and suppression of mitochondrial pathways, consistent with its known vulnerability during early AD pathogenesis (19,20,22,23,97). In contrast, the choroid plexus displayed earlier immune-associated transcriptional changes, supporting emerging evidence that barrier-associated neuroimmune interfaces undergo substantial inflammatory remodeling during aging (19,20,26,27). These observations reinforce the concept that intrinsic regional properties strongly influence susceptibility to age-associated molecular stress.

Sex emerged as a major modifier of aging trajectories. Females consistently exhibited stronger immune pathway enrichment and greater suppression of mitochondrial metabolic pathways than males, despite males showing transiently higher DEG burdens at intermediate aging stages. Similarly, in the independent AD cohort, males demonstrated broader transcriptional perturbation whereas females exhibited more coordinated enrichment of inflammatory and metabolic pathways. These findings suggest that pathway-level organization may provide greater biological insight into disease vulnerability than DEG burden alone and align with growing evidence that sex-specific differences in immune regulation, metabolism, and mitochondrial biology contribute to AD susceptibility (69,71,72,75,76,98–100).

Notably, the molecular alterations identified in AD closely mirrored those observed during physiological aging. Similar inflammatory and mitochondrial pathways were progressively altered during aging and became substantially amplified in AD, supporting a continuum model in which aging establishes a vulnerable immune– metabolic state that predisposes the brain to neurodegenerative pathology (52,84,101). Whereas physiological aging was associated predominantly with gradual structural remodeling and metabolic adaptation, AD was characterized by extensive neuroimmune activation together with pronounced suppression of oxidative metabolism. Together, these observations suggest that AD may emerge through amplification of aging-associated immune and metabolic dysfunction rather than through entirely distinct molecular mechanisms.

The principal novelty of this work lies in integrating three dimensions that are frequently investigated independently: regional aging heterogeneity, sex-dependent vulnerability, and mitochondrial–immune coupling. By combining mouse aging atlas (19) with independent human AD cohorts (52,74), our analyses support a unified model in which normal aging establishes a regionally patterned immune–metabolic state, females show stronger engagement of this state, and AD represents amplification of these aging-associated molecular programs. Positioning mitochondrial metabolism centrally within this framework provides a mechanistic perspective that extends beyond traditional amyloid-centric models (30–32,83,86,87) and highlights bioenergetic decline as a major component of aging-associated disease susceptibility.

Several limitations should be acknowledged. First, the analyses were performed on bulk transcriptomic datasets and therefore do not fully resolve cell-type-specific contributions. Given the central role of microglia, astrocytes, oligodendrocytes, vascular cells, and neuronal subtypes in aging and AD, future studies using single - cell and spatial transcriptomic approaches will be important to refine these observations (18,20,91,96,97,102– 105). Second, cross-species integration introduces variability related to species differences, regional sampling, and experimental design, although the conservation of major signatures across datasets supports the robustness of the findings. Third, our study establishes association rather than mechanistic causation between immune activation and mitochondrial dysfunction; experimental studies will be required to determine the temporal ordering and functional interdependence of these pathways. Finally, sex differences observed in transcriptomic profiles may also be influenced by hormonal status, environmental exposures, and systemic metabolic factors not fully captured within publicly available datasets (55,64–69).

In summary, our findings support a model in which coordinated immune activation and mitochondrial metabolic decline progressively reshape the aging brain, creating a molecular environment permissive for neurodegeneration. By integrating regional aging signatures, sex-dependent vulnerability, and conserved AD-associated pathways, this study identifies mitochondrial–immune coupling as a reproducible feature linking physiological aging to Alzheimer’s disease and highlights mitochondrial-immune interactions as a promising therapeutic axis for age-associated neurodegeneration (88,90,92–94).

## METHODS

### Lead contact

Further information and requests for resources, data and code availability should be directed to Joseph W. Guarnieri and Amita Pal.

### Materials availability

This study did not generate new, unique reagents or materials. All datasets analyzed in this research were obtained from publicly accessible datasets. The specific references and dataset accession numbers are listed in the “Experimental Model and Subject Details” below. Custom pathways used for gene set analysis are described in detail in “Methods Details” below.

### Data and code availability

All datasets analyzed in this study are publicly available through the Gene Expression omnibus under the accessions listed above (GSE212336, GSE104704, GSE125583). Code and gene lists utilized in this paper, can be found at https://github.com/shehbeel. Any additional information required to reanalyze the data reported in this paper is available from the lead contact upon request.

### Experimental Model and Subject Details

Publicly available transcriptomic datasets were obtained from the Gene Expression Omnibus and comprised: (i) a mouse brain aging atlas spanning multiple brain regions, ages, and sexes [GEO: GSE212336] (19); (ii) a human aging-to-AD transition cohort profiling lateral temporal cortex tissue from healthy young adults, healthy aged adults, and AD donors [GEO: GSE104704] (14); and (iii) an independent human AD validation cohort derived from fusiform gyrus tissue stratified by sex [GEO: GSE125583] (52). For the mouse aging dataset (GSE212336), the entorhinal cortex (EC) and choroid plexus (CP) were selected for in-depth analysis based on their established relevance to AD pathology and neuroimmune surveillance, respectively. 3-month-old mice served as the young reference baseline, and aging-associated comparisons were performed at 12, 15, 18, and 21 months in both male and female animals. Sample sizes ranged from n = 3–6 biological replicates per sex, age, and brain-region group. For the human aging-to-AD cohort (GSE104704), comparisons were performed among young healthy controls, aged healthy controls, and aged Alzheimer’s disease samples, with n = 8–12 subjects per group. For the independent AD validation cohort (GSE125583), comparisons were performed between Alzheimer’s disease and control samples in sex-stratified analyses, The study included female controls (n = 33), male controls (n = 37), female AD subjects (n = 97), and male AD subjects (n = 121). Samples lacking key metadata required for the relevant analysis (age, sex, brain region, or disease status) were excluded prior to differential expression testing.

### Method Details

#### RNA-seq processing and normalization

Gene-level count matrices were imported into R (v4.3.1) and processed using the *DESeq2* package (v1.40.2) (106). Lowly expressed genes were removed prior to analysis by retaining only those with at least 10 raw counts in a minimum number of samples equal to the smallest experimental group, thereby reducing noise and improving dispersion estimation. Library size normalization was performed using the median-of-ratios method implemented in *DESeq2*, and a variance-stabilizing transformation was applied to normalized counts for visualization, heatmaps, and dimensionality reduction. Sample-level quality control was assessed by inspection of library sizes, per-sample dispersion estimates and Cook’s distance for outlier detection. Sample structure was further evaluated using principal component analysis on VST-transformed data, and potential batch effects associated with sequencing run, processed date or donor cohort were inspected visually and statistically; where confounding was detected and metadata permitted, the corresponding covariate was incorporated into the *DESeq2* design matrix.

#### Differential gene expression analysis

Differential gene expression analysis was performed using *DESeq2* using the Wald test on negative binomial generalized linear models (106). For the mouse aging dataset, all comparisons were conducted relative to the young 3-month control group, enabling direct quantification of transcriptional changes associated with aging; gene expression at each older time point was compared against this 3-month baseline, and analyses were stratified by brain region and sex to capture region-specific and sex-dependent transcriptional responses. For the human datasets, differential expression analyses was performed between AD and control samples, both in pooled and sex-stratified analyses designs. Genes were considered differentially expressed (DEGs) when their Benjamini–Hochberg adjusted *P* value below 0.05; effect sizes are reported as log2 fold change throughout. Independent filtering as implemented in DESeq2 was used to optimize multiple-test corrections.

#### Functional enrichment and gene set enrichment analyses

Over-representation analysis was conducted using the *clusterProfiler* package, focusing on Gene Ontology (GO) Biological Process, Cellular Component, and Molecular Function terms. Up- and down-regulated DEGs were analyzed separately to preserve directionality, with all expressed genes used as the statistical background. GO term redundancy was reduced using the simplify function with a semantic-similarity cutoff of 0.7, and enriched terms with Benjamini–Hochberg adjusted P < 0.05 were retained. In addition to GO collections, manually curated gene sets covering immune activation, inflammation, complement signaling, oxidative phosphorylation, mitochondrial metabolism, ATP synthesis, respiratory electron transport chain assembly, mitochondrial translation, integrated stress and unfolded-protein responses, mitophagy, RAAS signaling, PANoptosis/senescence and extracellular matrix organization were assembled from prior literature and assayed by Gene set enrichment analysis (GSEA) to support the integrated pathway-summary heatmaps (107). These customs are mitochondrial, metabolic, and immune gene sets as previously described (107–111). GSEA was performed on ranked gene lists generated from the DESeq2 Wald statistic (or shrunken log2 fold change where indicated) using the fgsea algorithm implemented in clusterProfiler (112). Enrichment was quantified using normalized enrichment scores (NES), with statistical significance defined as a false discovery rate (FDR) < 0.25.

#### Cross-dataset integration

Cross-dataset integration was performed by comparing the directional consistency of gene-level and pathway-level changes across the aging, human aging-to-AD, and independent human AD validation datasets. Because these datasets differed in species, sampled brain region and cohort design, integration emphasized convergence of biological pathways rather than one-to-one equivalence of individual DEGs. Pathway-level integration used signed NES values from GSEA to assess shared directionality across cohorts. Conserved signatures were defined as pathways or genes that showed consistent directionality in at least two independent datasets, with particular attention to coordinated immune activation and suppression of mitochondrial metabolism. Mouse gene symbols were mapped to their human orthologs using the Mouse Genome Informatics (MGI) and Ensembl biomaRt resources prior to cross-species comparison (113).

#### Visualization

Volcano plots were generated using the EnhancedVolcano package (v1.14.0), with log2 fold change plotted against -log10 adjusted *P* value; significance thresholds and labeled gene sets are specified in the corresponding figure legends. Bar plots, dot plots, lollipop plots and pathway-level summaries were produced with *ggplot2* (v3.5.0) (114). Heatmaps were generated using *pheatmap* and *ComplexHeatmap* (115). Brain-region schematics and anatomical illustrations were created in BioRender.com. Final figure panels were assembled in vector-based graphics software and arrangedaccording to the layout shown in the corresponding figure legends.

### Quantification and Statistical Analysis

All statistical analyses were performed in R (v4.3.1). Unless otherwise indicated, *P* values were adjusted for multiple testing using the Benjamin-Hochberg false-discovery rate procedure, and statistical significance was defined as adjusted *P* < 0.05. No statistical method was used to predetermine sample size; sample sizes reflect those of the original publicly available datasets. No data were excluded from the analyses except for the metadata-driven exclusions described above. The investigators were not blinded to group allocation, because group assignments (age, sex, disease status) were intrinsic properties of the publicly available samples and could not be masked during computational analyses.

## Author Contributions

Conceptualization: J.W.G., A.P.; Methodology: J.W.G, A.P.; Formal Analysis: J.W.G., A.P., S.A.; Investigation: J.W.G., A.P.; Data Curation: J.W.G., A.P.; Resources: J.W.G., A.P.; Writing – Original Draft: A.P., J.W.G.; Writing – Review & Editing: A.P., S.A., I.K., E.W., J.W.G.,; Visualization: J.W.G., A.P.; Funding Acquisition: J.W.G.; Supervision: J.W.G.

## Graphical abstract

**Graphical Summary:**
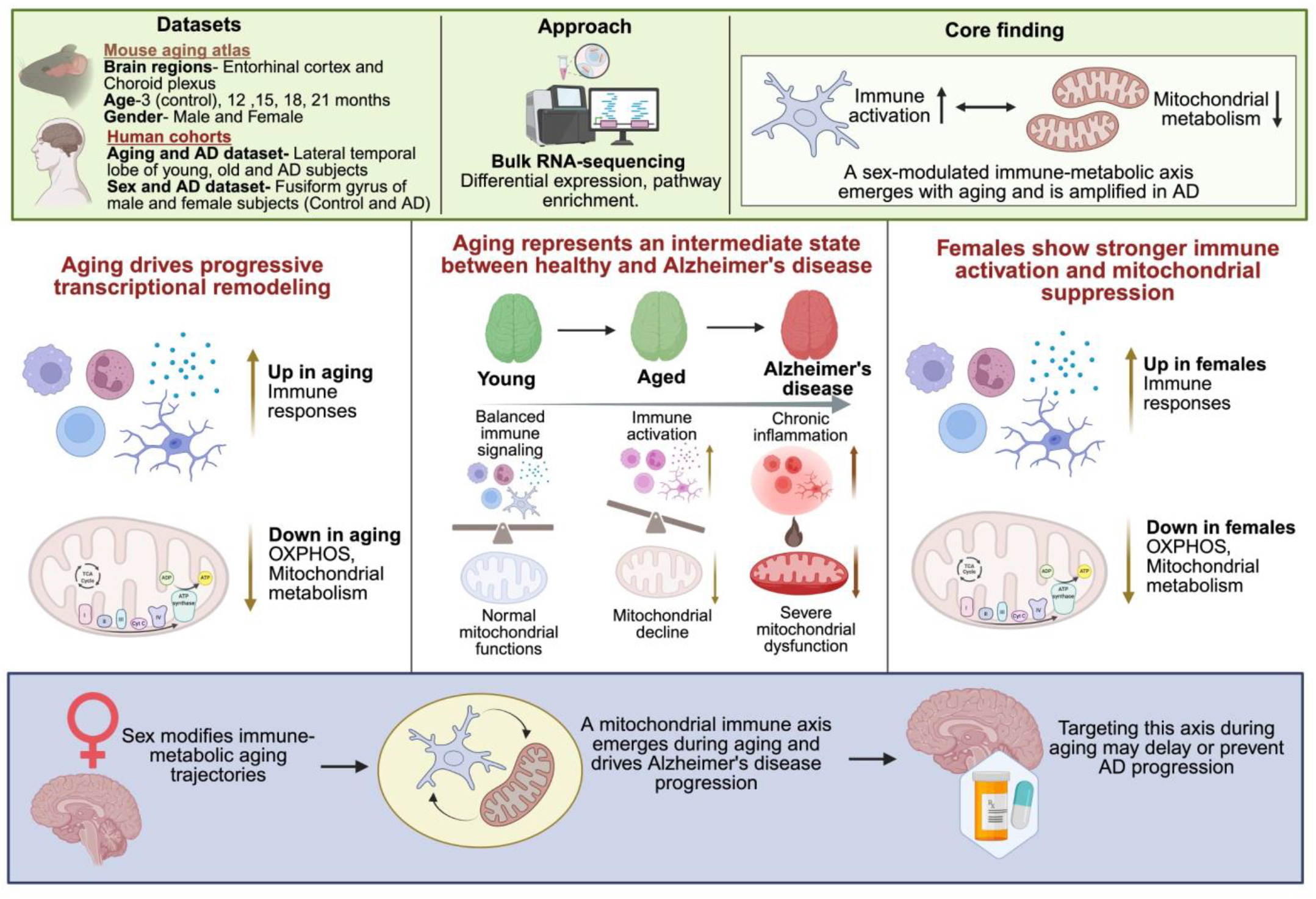
Integrative transcriptomics identifies a conserved mitochondrial–immune axis linking brain aging and Alzheimer’s disease: Integrative transcriptomic analyses of mouse brain aging and human aging-to-Alzheimer’s disease (AD) cohorts identify coordinated remodeling of mitochondrial, immune, and stress-response pathways during aging and neurodegeneration. Aging is associated with progressive alterations in oxidative phosphorylation, mitochondrial metabolism, mito-innate immune signaling, inflammatory pathways, and cellular stress-response programs, including integrated stress response (ISR), unfolded protein response (UPR), autophagy, and mitophagy. Comparative analyses demonstrate that female aging brains exhibit stronger immune activation and mitochondrial stress-associated transcriptional signatures relative to males. Aging-associated molecular signatures further overlap substantially with AD-related transcriptional programs, including enhanced inflammatory signaling, extracellular matrix remodeling, and inflammatory cell-death-associated pathways. Collectively, these findings support a model in which mitochondrial dysfunction promotes chronic neuroimmune activation and establishes a sex-dependent mitochondrial–immune vulnerability state that is further amplified during AD progression.

