## Supplementary Data for "A mitochondrial-immune axis drives the transcriptomic transition from brain aging to Alzheimer’s disease"

### Table of Contents

**Figure S1. Expanded region- and sex-stratified aging transcriptomic analyses**…...…………...………..2-3

**Figure S2. Expanded transcriptional trajectory analyses across the aging-to-Alzheimer's disease continuum**……..………………………………………………………………………...……………………….4-5

**Figure S3. Expanded validation analyses of immune and mitochondrial signatures in an independent Alzheimer's disease cohort**………………………………..…………………………………………………….5-6

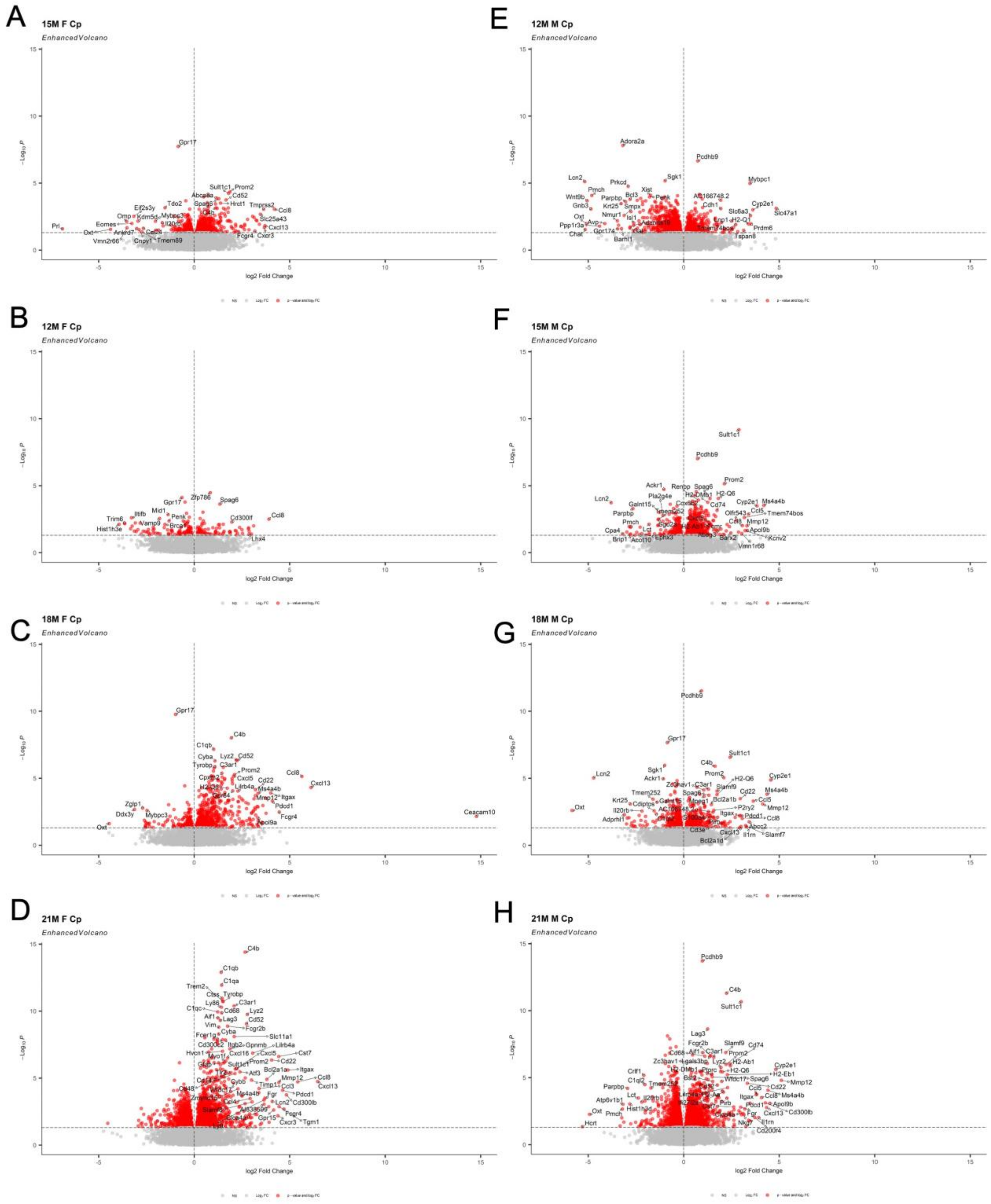


F

**Figure S1. Expanded region- and sex-stratified aging transcriptomic analyses.** Enhanced volcano plots of all age-, sex- and region-specific comparisons relative to 3-month young controls, supporting the summarized DEG counts in Fig. 1B- D. (A) Female (left) and male (right) volcano plots for the CP at 12, 15, 18 and 21 months. (B) Female (left) and male (right) volcano plots for the EC at the same time points. Each point represents one gene plotted by $\log_2$ fold change against $-\log_{10}$(adjusted $P$). Significant genes (adjusted $P < 0.05$) are shown in red; non-significant genes in grey. Selected top genes per comparison are labeled. Late-stage comparisons show broader transcriptional dispersion and pronounced upregulation of immune and stress-response genes.

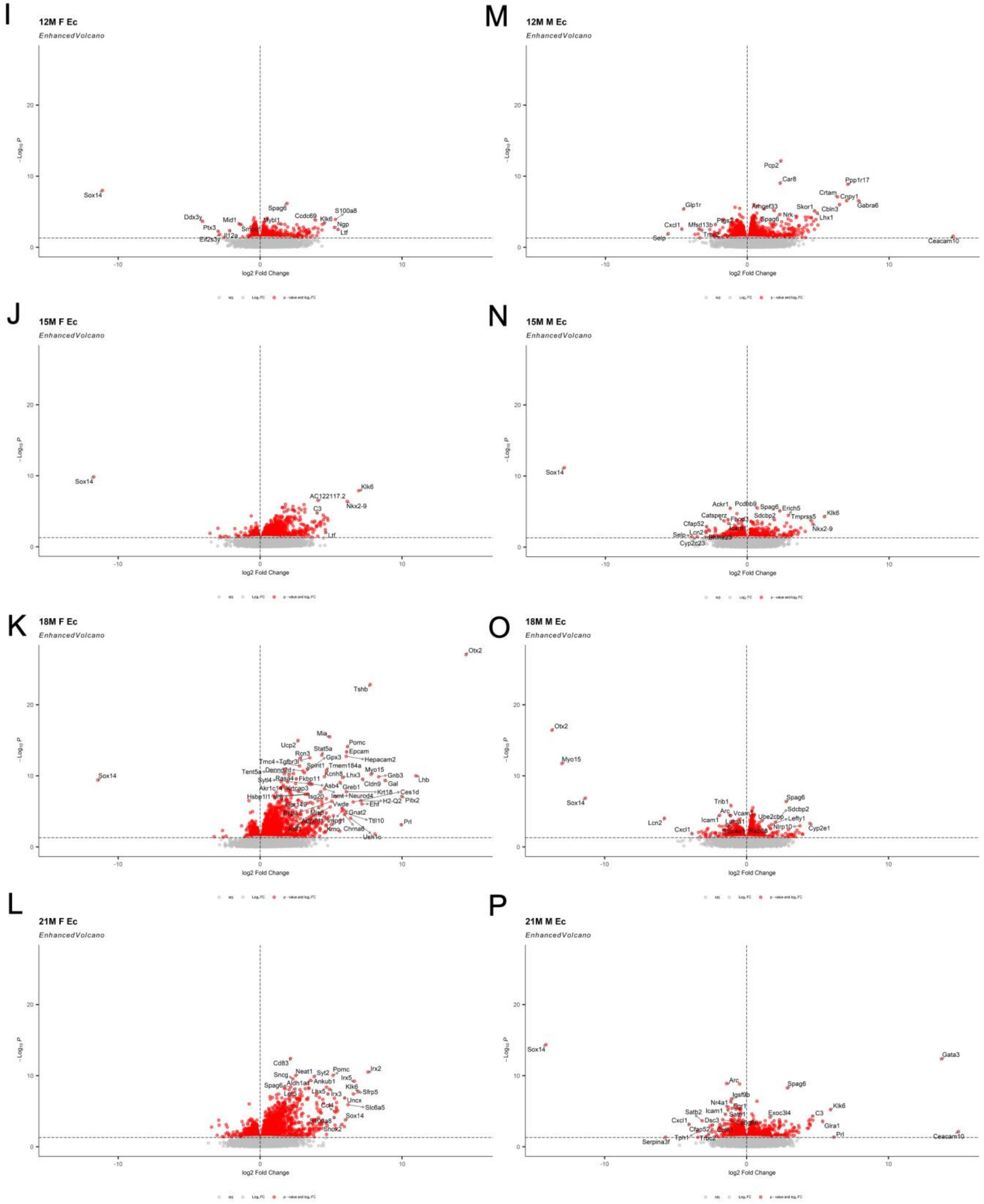


**Figure S1. Expanded region- and sex-stratified aging transcriptomic analyses (continued).**

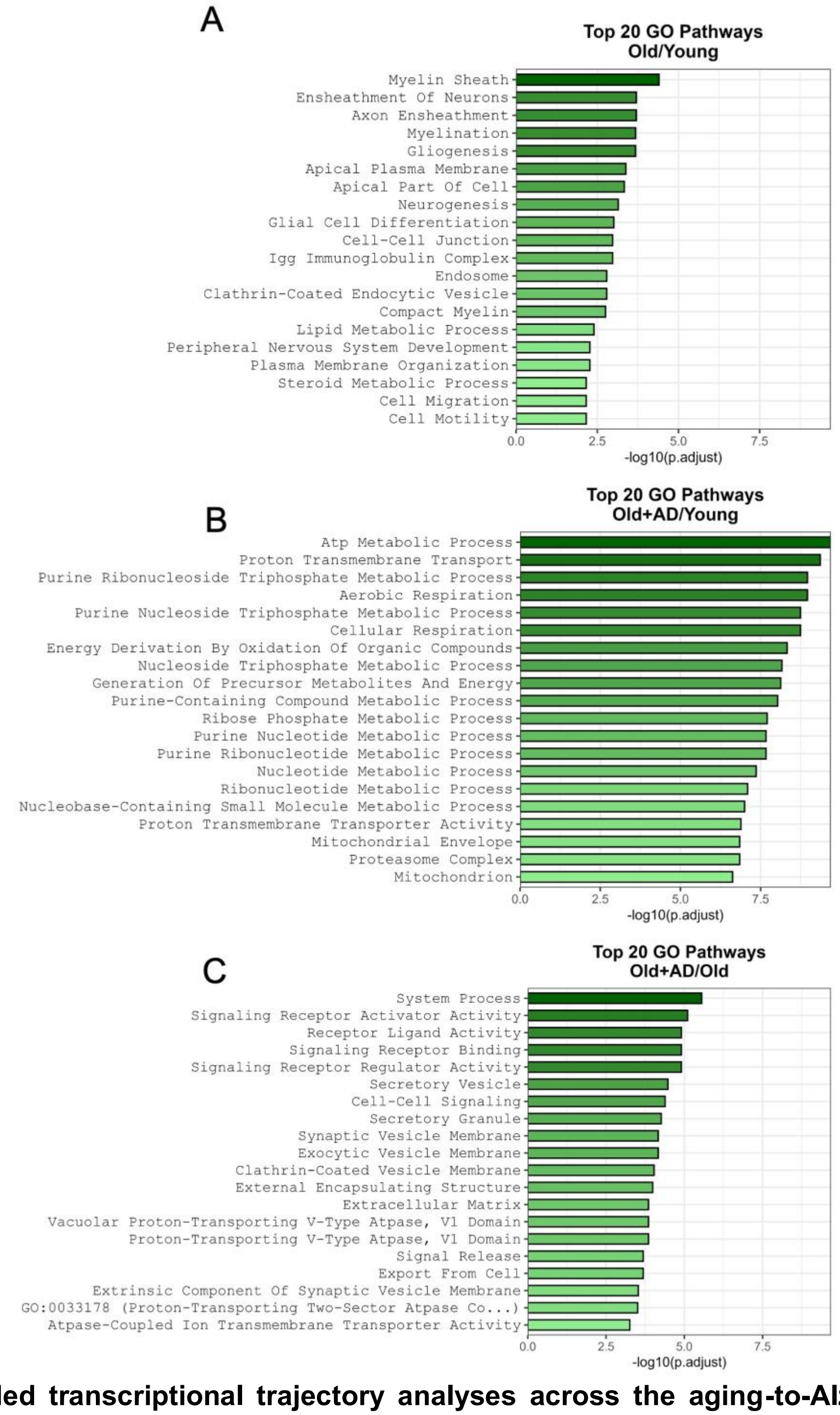


**Figure S2. Expanded transcriptional trajectory analyses across the aging-to-Alzheimer's disease continuum.** (A) Bar plots of the top 20 enriched GO Biological Process terms for each comparison along the aging-to-AD continuum (Old/Young, Old+AD/Young and Old+AD/Old). Bar length represents $-\log_{10}$(adjusted *P*). Physiological aging is dominated by myelination, gliogenesis and structural remodeling pathways; AD-associated comparisons are dominated by mitochondrial metabolic processes, ATP metabolism, aerobic respiration and proton transmembrane transport. (B) Enhanced volcano plots for the same three comparisons. Each point represents one gene plotted by $\log_2$ fold change against $-\log_{10}$(adjusted *P*). Significant genes (adjusted $P < 0.05$) are highlighted in red; non-significant genes in grey. AD-associated comparisons show broader transcriptional dispersion and stronger suppression of oxidative-metabolism-associated genes than physiological aging.

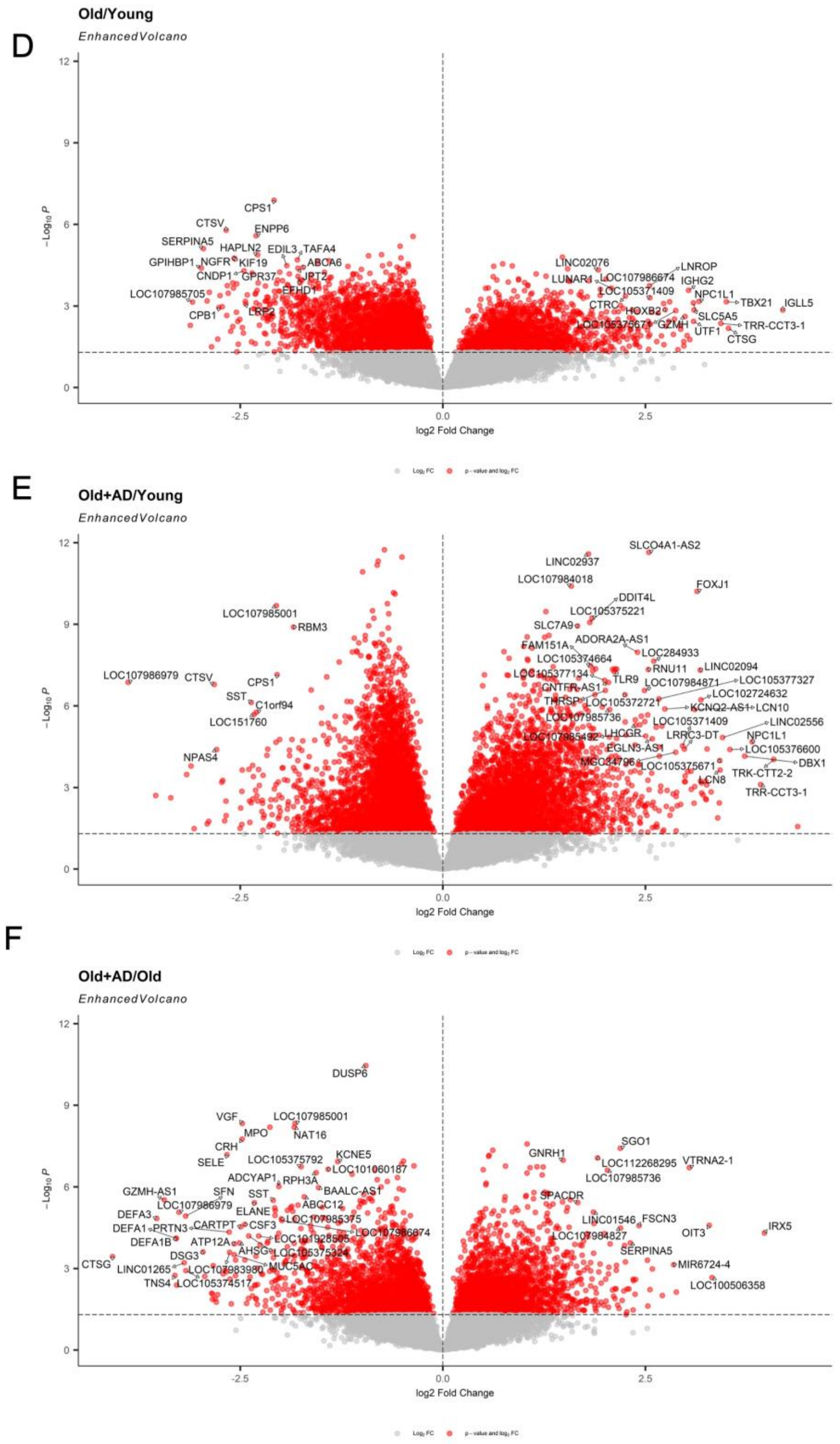


**Figure S2. Expanded transcriptional trajectory analyses across the aging-to-Alzheimer's disease continuum (continued).**

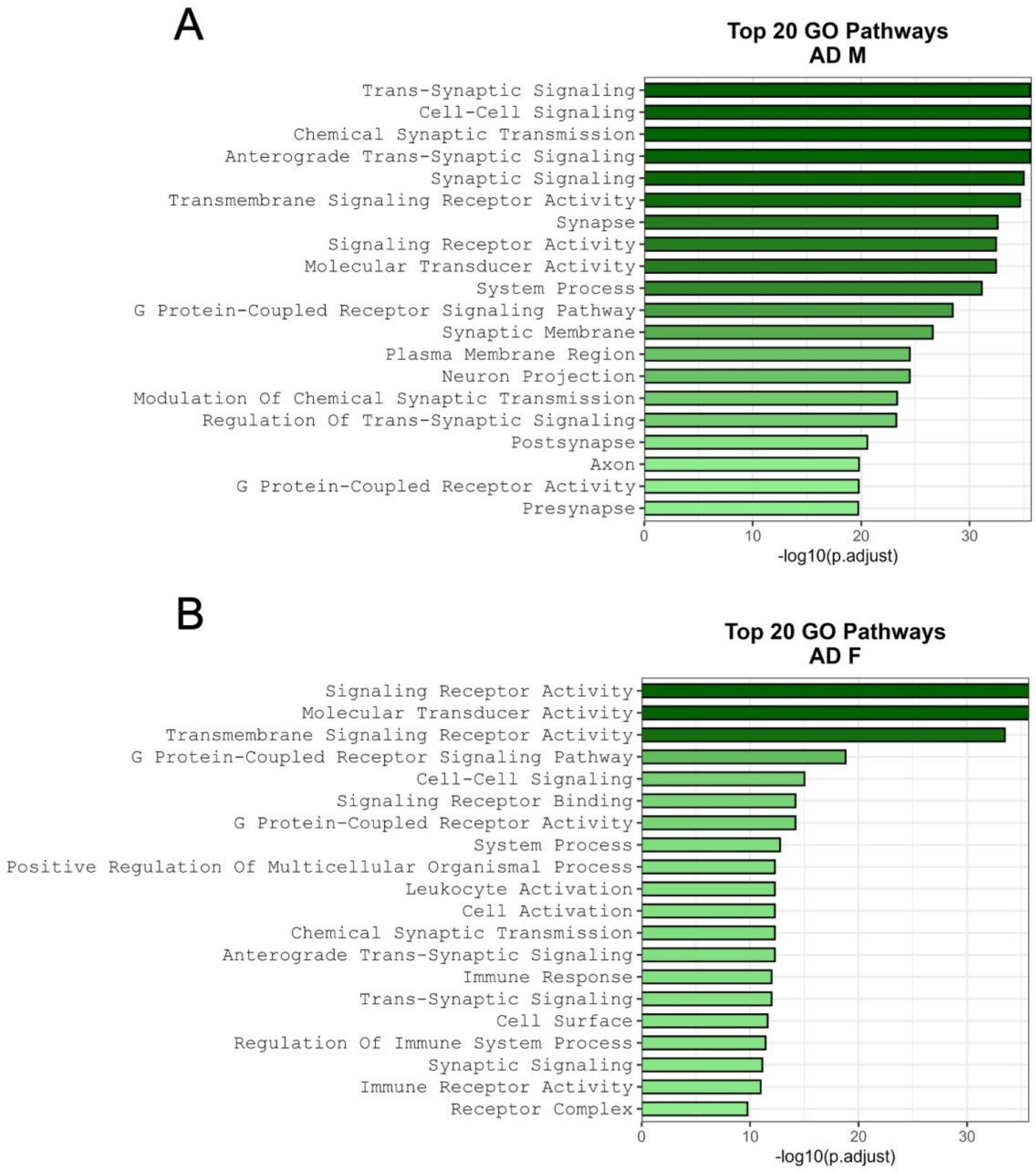


**Figure S3. Expanded validation analyses of immune and mitochondrial signatures in an independent Alzheimer's disease cohort.** (A) Bar plots of the top 20 enriched GO Biological Process terms for the AD M and AD F comparisons. Bar length represents $-\log_{10}$(adjusted *P*). AD samples of both sexes show enrichment of trans-synaptic signaling, receptor activity and synaptic communication pathways together with increasing immune-regulatory and leukocyte-associated programs. (B) Enhanced volcano plots for the AD M and AD F comparisons spanning the full transcriptome. Each point represents one gene plotted by $\log_2$ fold change against $-\log_{10}$(adjusted *P*). Significant genes (adjusted $P < 0.05$) are highlighted in red; non-significant genes in grey. Selected top genes are labeled. AD samples show coordinated suppression of mitochondrial respiratory-chain genes together with induction of inflammatory transcripts.

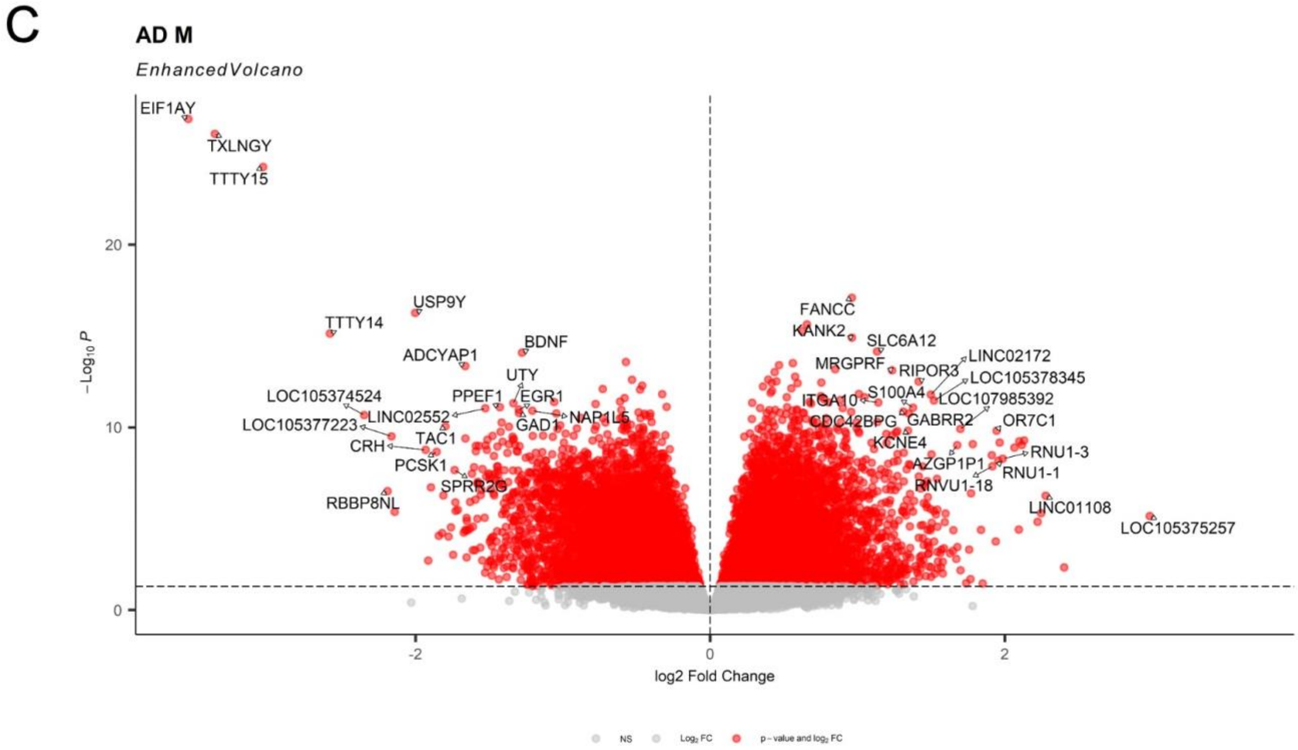


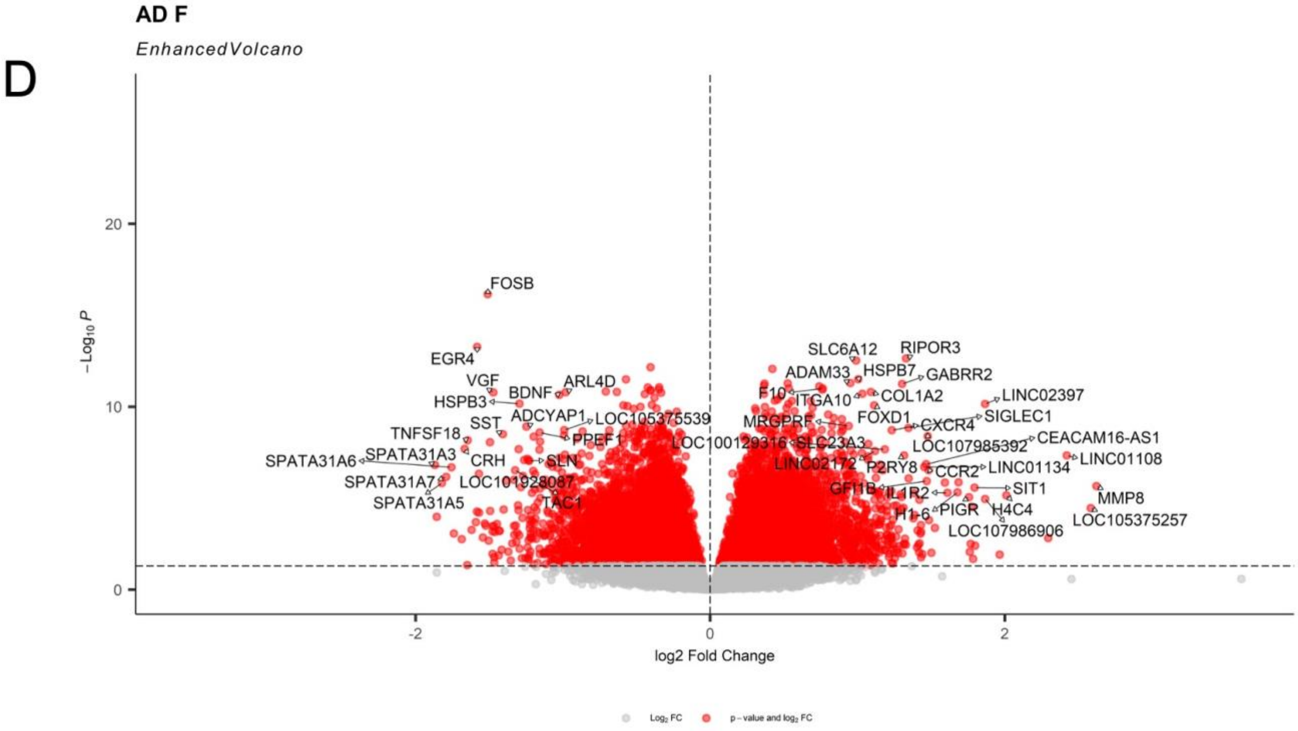


**Figure S3. Expanded validation analyses of immune and mitochondrial signatures in an independent Alzheimer's disease cohort (continued).**
